# Hyperlocal Variation in Soil Iron and Rhizosphere Microbiome Determines Disease Development in Amenity Turfgrass

**DOI:** 10.1101/2020.08.17.255265

**Authors:** Ming-Yi Chou, Smita Shrestha, Renee Rioux, Paul Koch

## Abstract

Dollar spot, caused by the fungal pathogen *Clarireedia spp*., is an economically important disease of amenity turfgrass in temperate climates worldwide. This disease often occurs in a highly variable manner, even on a local scale with relatively uniform environmental conditions. The objective of this study was to investigate mechanisms behind this local variation, focusing on contributions of the soil and rhizosphere microbiome. Turfgrass, rhizosphere, and bulk soil samples were taken from within a 256 m^2^ area of healthy turfgrass, transported to a controlled environment chamber, and inoculated with *C. jacksonii*. Bacterial communities were profiled targeting the 16s rRNA gene, and 16 different soil chemical properties were assessed. Despite their initial uniform appearance, the samples differentiated into highly susceptible and moderately susceptible groups following inoculation in the controlled environment chamber. The highly susceptible samples harbored a unique rhizosphere microbiome with lower relative abundance of antibiotic-producing bacterial taxa and higher predicted abundance of genes associated with xenobiotic biodegradation pathways. In addition, stepwise regression revealed that bulk soil iron content was the only significant soil characteristic that positively regressed with decreased dollar spot susceptibility during the peak disease development stage. These findings suggest that localized variation in soil iron induces the plant to select for a particular rhizosphere microbiome that alters the disease outcome. More broadly, further research in this area may indicate how plot-scale variability in soil properties can drive variable plant disease development through alterations in the rhizosphere microbiome.

**IMPORTANCE:** Dollar spot is the most economically important disease of amenity turfgrass, and more fungicides are applied targeting dollar spot than any other turfgrass disease. Dollar spot symptoms are small (3-5 cm), circular patches that develop in a highly variable manner within plot-scale even under seemingly uniform conditions. The mechanism behind this variable development is unknown. This study observed that differences in dollar spot development over a 256 m^2^ area were associated with differences in bulk soil iron concentration and correlated with a particular rhizosphere microbiome. These findings provide important clues for understanding the mechanisms behind the highly variable development of dollar spot, which may offer important clues for innovative control strategies. Additionally, these results also suggest that small changes in soil properties can alter plant activity and hence the plant-associated microbial community which has important implications for a broad array of important agricultural and horticultural plant pathosystems.

## INTRODUCTION

Dollar spot on cool-season turfgrasses in North America is caused by the fungus *Clarireedia jacksonii* and is the most economically important disease of amenity turfgrass in temperate climates around the world (1). It causes roughly circular patches of bleached turfgrass 3 to 5 cm in diameter that can blight the stand and reduce the functionality of the site for recreational purposes (2). The primary host of dollar spot is creeping bentgrass (*Agrostis stolonifera*), and a lack of host resistance or effective cultural control strategies has made dollar spot the target of more fungicide applications than any other turfgrass disease (3). Heavy reliance on synthetic fungicides has led to the development of fungicide resistant fungal populations (4), imposes a significant financial burden on the turfgrass manager (5), and increases the risk of human and environmental contamination resulting from repeated chemical exposures (6). The development of dollar spot symptoms is often highly variable within several meters distance, even in uniformly managed turfgrass with nearly identical environmental conditions (7). While it is not known why dollar spot symptoms develop in such a variable manner, one plausible explanation is a link to hyperlocal variations in microbial antagonists or variations in soil physical, chemical, or biological properties.

Spatial variation in plant disease is often observed in both managed and natural plant systems, and most studies on variation in plant disease incidence and severity have been conducted in large-scale agricultural fields over tens or hundreds of hectares. A. Adiobo et al. (8) observed that the physicochemical and microbial properties of andosols suppressed *Pythium myriotylum* root rot in cocoyam (*Xanthosoma sagittifolium*) more effectively than ferralsols. Varied susceptibility to disease in adjacent fields with similar soil physicochemical characteristics has commonly been attributed to disease suppressive or disease conducive soils and is often influenced by cropping history (9, 10). Though on a larger scale than the variation observed in dollar spot, the pathogen suppression function of a specific suppressive soil has provided some clues as to how the same soil type could have dramatically different pathogen suppression functions. Enrichment of the antagonistic microbial population in the rhizosphere often serves as the key plant pathogen suppression mechanism in previously characterized disease suppressive soils (11). As a classic example, enriched antibiotic 2,4-diacetylphloroglucinol-producing fluorescent *Pseudomonas* species led to a reduction in take-all disease when found in the rhizosphere of wheat and flax (12).

The rhizosphere microbiome and its functions are co-determined by both the plant and the soil. The host plant produces root exudates that recruit particular microbes from within the soil (11). The soil harbors varied microbial communities shaped by soil type and associated properties, such as structure and pH (13). Therefore, the rhizosphere microbiome and its microbial disease suppressive function can shift following changes in the soil environment. H. Peng et al. (14) varied the chemical and physical properties of *Fusarium oxysporum* f. sp. *cubense* suppressive and conducive soils and showed that soil physicochemical traits can mediate suppressiveness of both suppressive and conducive soils against the pathogen’s chlamydospores. This suggests that soil physicochemical and microbial properties can cooperatively affect plant disease suppression in agricultural fields.

Soil spatial variation in microbial properties is often studied at multiple levels, including micro, plot, field, landscape and regional scales (15, 16). Over a small plot-scale, spatial variation of smut disease (*Ustilago syntherismae*) on crabgrass (*Digitaria sanguinalis*) was influenced by both pathogen spore density and spatial location (17). However, soil property influences were not investigated in this study and spores or other long-distance dispersal mechanisms have never been associated with dollar spot in a field environment (2). High spatial variations in soil physicochemical and microbial properties were observed in a managed grassland, including a wide range of soil pH, nitrogen content, microbial biomass, and microbial catabolism profiles within the scales of several centimeters to meters (18), but the impact of these variations on plant-pathogen interactions remained unclear. Recently, Z. Wei et al. (19) examined disease variation in tomato (*Solanum lycopersicum*) and observed that the rhizosphere soil bacterial community effectively predicted the severity of the soil-borne bacterial disease *Ralstonia solanacearum*. Similarly, S. Chen et al. (20) observed differences in rhizosphere bacterial community structure, diversity, acid phosphatase activity, root iron content, and bulk soil calcium and magnesium between healthy and unhealthy blueberry plants (*Vaccinium corymbosum*). These results again indicate the importance of both soil chemical properties and the rhizosphere microbiome on plant health over a field-scale or smaller. However, it remains unknown whether the rhizosphere and/or bulk soil microbiome impacts the disease severity of a foliar fungal pathogen when interacting with specific soil chemical properties.

In this study, various factors contributing to the localized variation in dollar spot development on monocultured turfgrass was studied. Rhizosphere and bulk soil microbiomes as well as soil chemical properties were examined to determine possible causes for the highly variable spatial nature of dollar spot development. We hypothesized that soil chemical properties and the rhizosphere microbiome are both significant variables in determining dollar spot disease susceptibility in a uniformly managed and monocultured turfgrass system. Turfgrass is an excellent system to study this phenomenon because the high plant density allows for robust sampling over a small scale. The initial 132 cm^2^ surface area turfgrass soil plug harbored an estimated 1,200 individual creeping bentgrass plants, and each sub-sample derived from the soil core contained 10 to 15 individual plants. By understanding the factors that drive variation in dollar spot disease development within a plot-scale in a high-density monoculture system, we may discover mechanisms that can be targeted for improved biological management of a number of important plant pathogens.

## RESULTS

### Dollar spot development

Dollar spot development was measured as decrease of greenness over time in order to standardize the quantification of disease symptoms as lesion shape and color can be difficult to determine with simple visual assessments. The resulting greenness decay curve followed a sigmoidal decay pattern (r=0.9286 and p-value<0.0001) (Fig.1). Disease symptoms initially developed within two days after inoculation (DAI), then increased rapidly over the next four to twelve DAI, before slowing during the saturation phase on 14 to 16 DAI. Substantial differences in symptom severity between samples started showing up on four DAI and differences remained apparent throughout the incubation.

**Figure 1.**
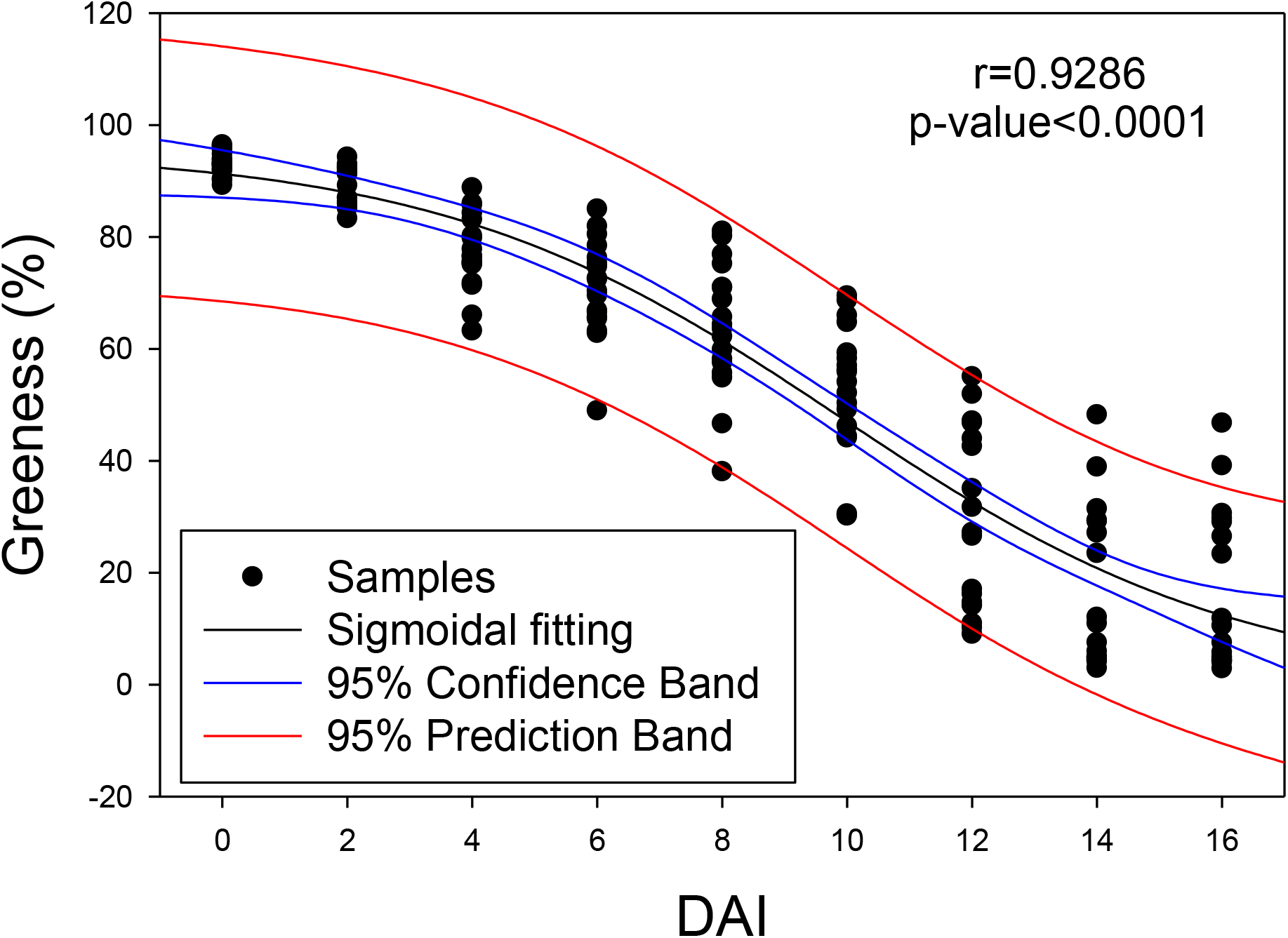
Dollar spot development as indicated by turf greenness decay curve fitted with sigmoidal model (r=0.9286, p<0.0001) throughout 16 days of incubation after dollar spot inoculation (n=18). DAI stands for days after inoculation with *C. jacksonii*.

### Attributing soil bacterial community difference as a function of disease variability

Turf samples were grouped into high, medium, and low disease according to the disease severity of each DAI. The bacterial microbiome from rhizosphere and bulk soil associated with each sample, which had been separated prior to inoculation, was then assessed to see if the microbiome structure explained turfgrass responses to *C. jacksonii* inoculation. The rhizosphere bacterial community differed between high and low disease severity groups when categorized based on severity between 4 and 10 DAI according to permutational analysis of variance (PERMANOVA) (Table 1). There were no differences in bacterial community structure found between high and low disease severity groups when categorized according to initial disease development (DAI 0-2) or the disease saturation phase (DAI 12-16). In addition, no differences in the bulk soil bacterial community were found among the disease severity groups throughout the entire incubation (Table 1). The period that the rhizosphere soil microbiome showed differences in structure between the high and low disease groups (4-10 DAI) matched the backslope of the disease development curve (Fig. 1), which suggested that the initial soil rhizosphere microbiome can affect the peak dollar spot development. The samples were then re-categorized according to their disease status during the peak disease development stage (4-10 DAI) to make the peak disease development period as the target of prediction instead of any single day within this period. The samples initially categorized as high disease during the 4 to 10 DAI period never shifted into the low severity group and vice versa, so the 18 samples naturally broke into two groups except for one sample that stayed in the medium disease group throughout the study and was excluded from further analysis. Further analyses were performed based on breaking the samples into nine highly susceptible (HS) samples and eight moderately susceptible (MS) samples.

**Table 1.** Paired-PERMANOVA analysis of turf-associated soil microbiome prior to *C. jacksonii* inoculation as categorized based on disease level (high, medium, and low) after inoculation of *C. jacksonii* and throughout the incubation. Asterix indicates the significance level: * p<0.05 and ** P<0.01. DAI stands for Days after inoculation of *C. jacksonii*.

| Pair-comparison |  | 0 DAI |  | 2 DAI |  | 4 DAI |  | 6 DAI |  | 8 DAI |  |
| --- | --- | --- | --- | --- | --- | --- | --- | --- | --- | --- | --- |
|  |  | R<sup>2</sup> | P-value | R<sup>2</sup> | P-value | R<sup>2</sup> | P-value | R<sup>2</sup> | P-value | R<sup>2</sup> | P-value |
| Rhizosphere | High vs Low | 0.116 | 0.126 | 0.124 | 0.090 | 0.138 | 0.006** | 0.139 | 0.003** | 0.118 | 0.021* |
|  | High vs Med | 0.080 | 1.000 | 0.100 | 0.603 | 0.136 | 0.012* | 0.112 | 0.090 | 0.103 | 0.345 |
|  | Low vs Med | 0.115 | 0.117 | 0.089 | 1.000 | 0.103 | 0.147 | 0.116 | 0.036* | 0.096 | 0.807 |
| Bulk | High vs Low | 0.125 | 0.144 | 0.101 | 0.597 | 0.087 | 1.000 | 0.089 | 1.000 | 0.085 | 1.000 |
|  | High vs Med | 0.094 | 0.978 | 0.096 | 0.939 | 0.086 | 1.000 | 0.112 | 0.372 | 0.122 | 0.207 |
|  | Low vs Med | 0.093 | 0.984 | 0.087 | 1.000 | 0.076 | 1.000 | 0.088 | 1.000 | 0.099 | 0.549 |
| Pair-comparison |  | 10 DAI |  | 12 DAI |  | 14 DAI |  | 16 DAI |  |  |  |
|  |  | R<sup>2</sup> | P-value | R<sup>2</sup> | P-value | R<sup>2</sup> | P-value | R<sup>2</sup> | P-value |  |  |
| Rhizosphere | High vs Low | 0.126 | 0.024* | 0.091 | 1.000 | 0.091 | 1.000 | 0.091 | 1.000 |  |  |
|  | High vs Med | 0.128 | 0.036* | 0.080 | 1.000 | 0.096 | 0.840 | 0.096 | 0.816 |  |  |
|  | Low vs Med | 0.109 | 0.096 | 0.092 | 1.000 | 0.100 | 0.540 | 0.100 | 0.573 |  |  |
| Bulk | High vs Low | 0.080 | 1.000 | 0.080 | 1.000 | 0.091 | 1.000 | 0.091 | 1.000 |  |  |
|  | High vs Med | 0.081 | 1.000 | 0.088 | 1.000 | 0.092 | 1.000 | 0.092 | 1.000 |  |  |
|  | Low vs Med | 0.080 | 1.000 | 0.076 | 1.000 | 0.068 | 1.000 | 0.068 | 1.000 |  |  |

### Comparison of rhizosphere bacterial communities of highly susceptible and moderately susceptible turfgrass

Two-dimensional principal coordinate analysis showed that distinct bacterial community structures existed between the bulk and rhizosphere soil and between the rhizosphere soil of HS and MS samples (Fig. 2). PERMANOVA statistically confirmed the visual observations of bacterial community composition differences between sample types (Fig. 2a) and susceptibility groups of rhizosphere soil (Fig. 2b). Although the overall rhizosphere bacterial compositions are different between MS and HS turfgrass, the major microbial taxa are identical when analyzed at family and genus levels with less than 20% and more than 75% of the taxa unidentified at each taxonomic level, respectively (Fig. 3). The dominant families identified included *Gemmataceae*, *Pirellulaceae*, *Chitinophagaceae*, *Pedospheraceae*, and *Burkholderiaceae* (Fig. 3a) and the dominant genus’ identified included *Flavobacterium*, *Haliangium*, *Chthoniobacter*, *Pirellula*, and *Gaiella* (Fig. 3b). The majority of the rhizosphere soil amplicon sequence variants (ASVs) are shared between the HS and MS turfgrass (8077) with more ASVs being unique to HS (1181) than MS (347) (Fig. 4). Highly susceptible turfgrass samples also had a higher species richness and β-diversity as shown using the Shannon index (Fig. 5).

**Figure 2.**
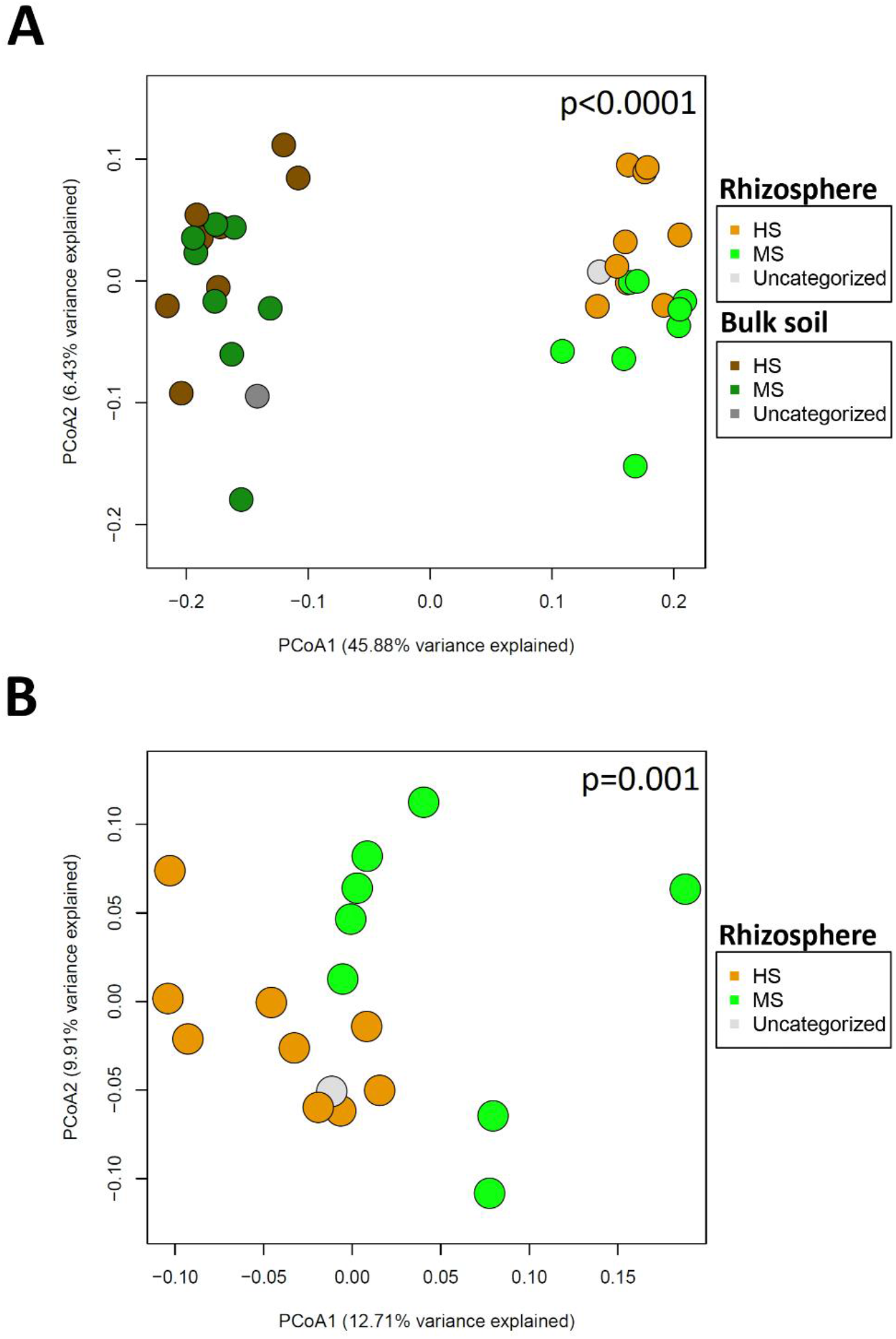
Principal coordinate analysis (PCoA) of bulk soil versus rhizosphere microbiome (a), and MS versus HS turfgrass rhizosphere microbiome (b). Significant differences between MS and HS samples were tested using PERMANOVA.

**Figure 3.**
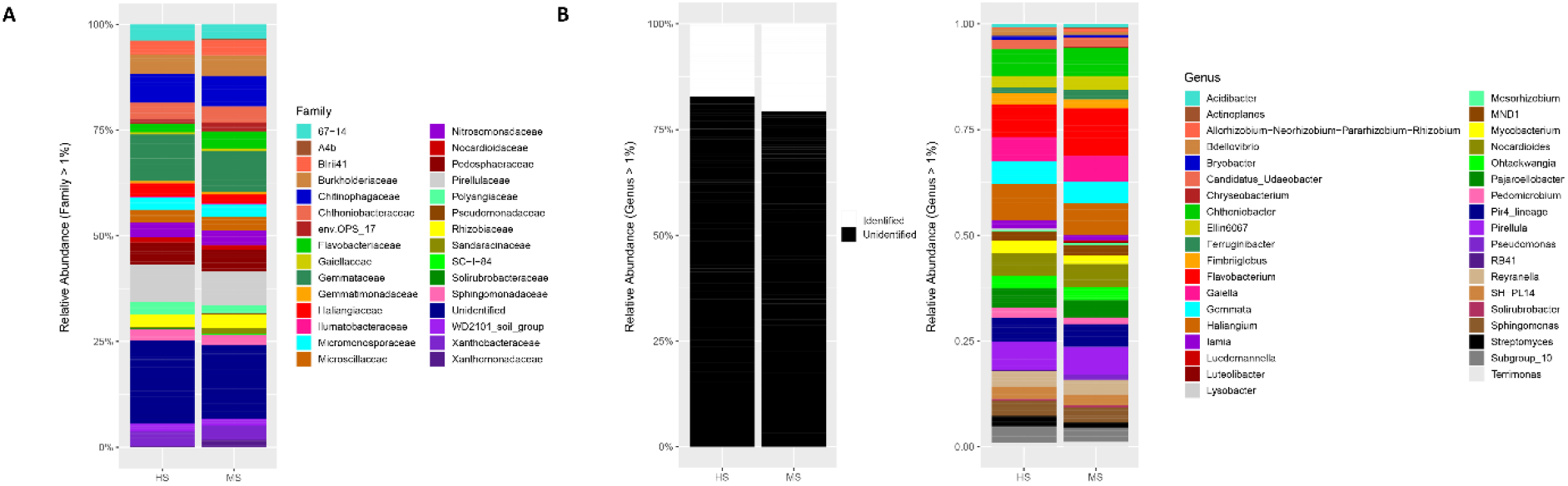
Relative abundance of rhizosphere microbiome from MS and HS turfgrass at Family (a) and Genus (b) level.

**Figure 4.**
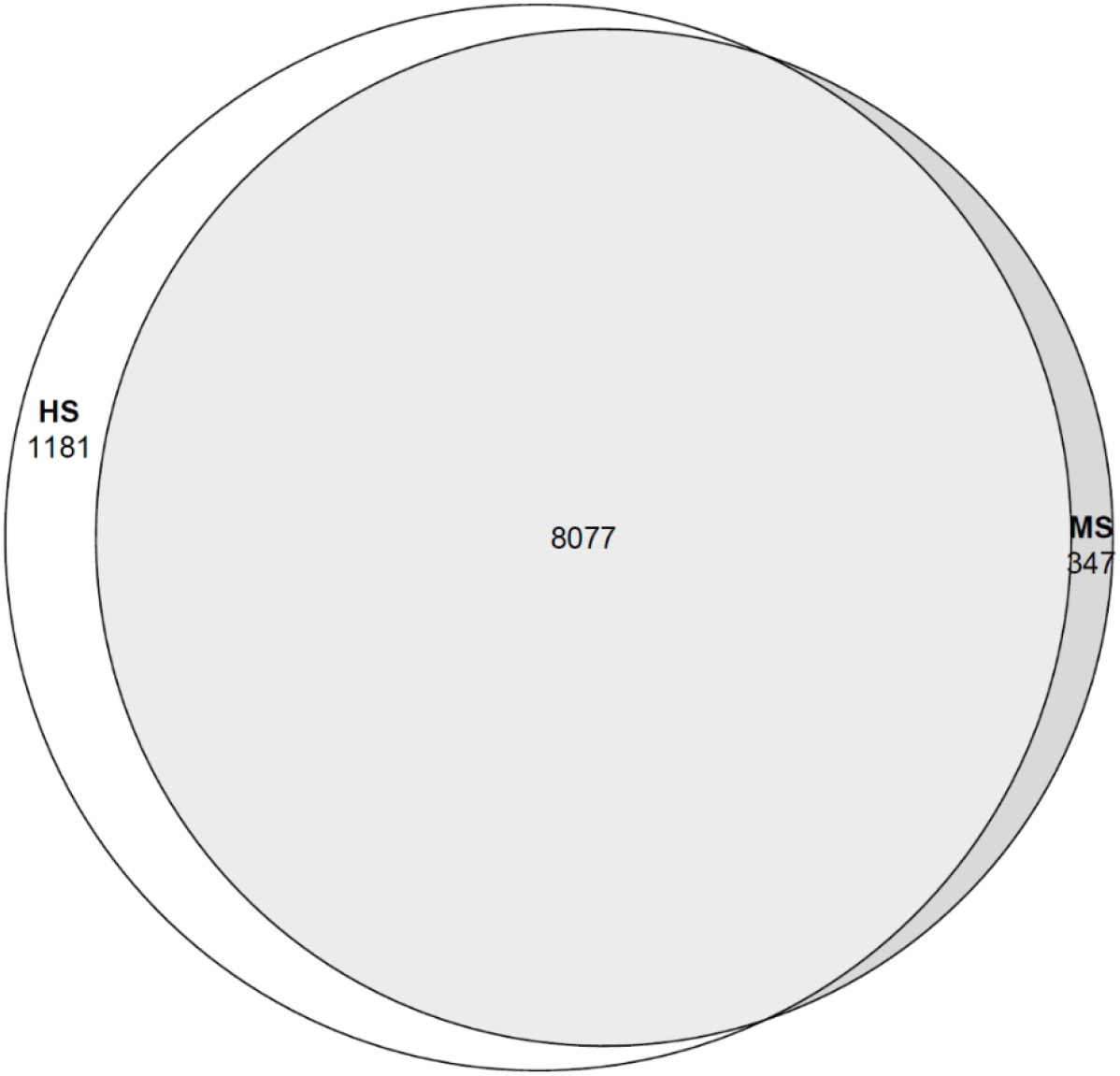
Proportions of shared and unique ASV between the MS and HS rhizosphere soil showed as Venn diagram.

**Figure 5.**
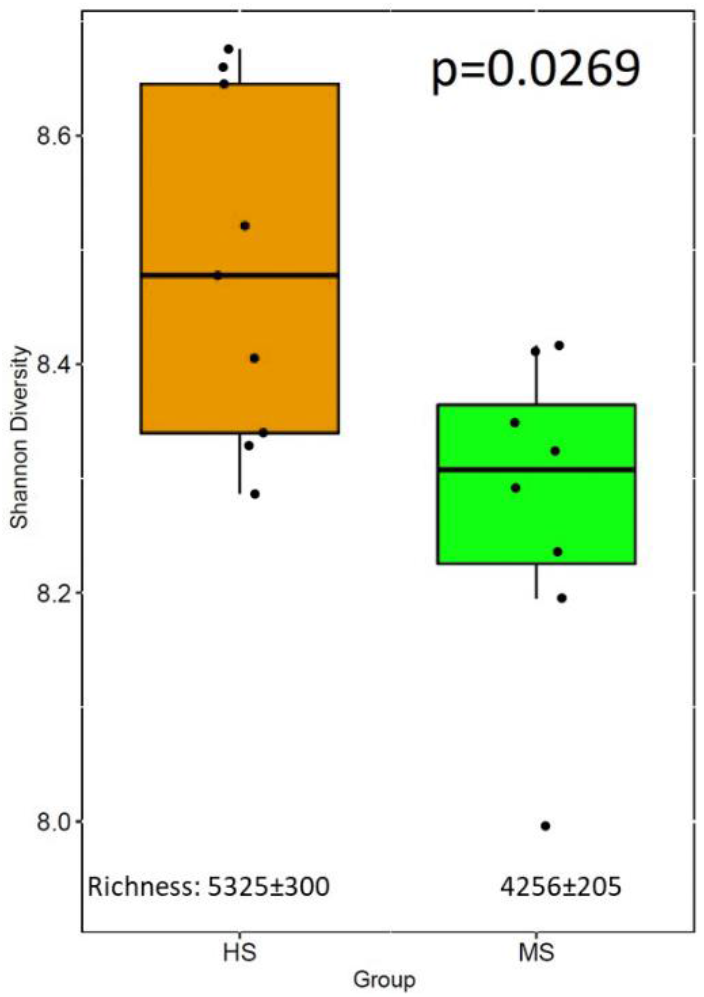
Bacterial Shannon diversity and richness of rhizosphere microbiome for turfgrass of MS and HS susceptibility group. The Shannon diversity significant difference was performed using a nonparametric Wilcoxon test.

In the rhizosphere, there were 28 families and 32 genera different in relative abundance between HS and MS samples according to Welch’s t-test (Fig. 6a). A balance analysis that accounted for the compositional nature of the dataset was also performed to detect the microbial signature for discerning the high and low disease rhizosphere bacterial community. The signatures were determined by searching the association between the factor for overall microbiome difference with the bacterial taxa balances defined as normalized log ratio of the geometric mean of the numerator and denominator bacterial taxa. The results showed that relative abundance log ratio of *Rhizobacter* (numerator) to *Microvirga* (denominator) at the genus level and *Solibacteraceae* subgroup3 (numerator) to *Saprospiraceae* (denominator) at the Family level were robust microbial signatures to differentiate the HS and MS turfgrass rhizosphere bacterial community with an adjusted area under the receiver operating characteristic curve for cross-validation equal to 0.9875 and 0.983 for genus and family level, respectively (Fig. 6b).

**Figure 6.**
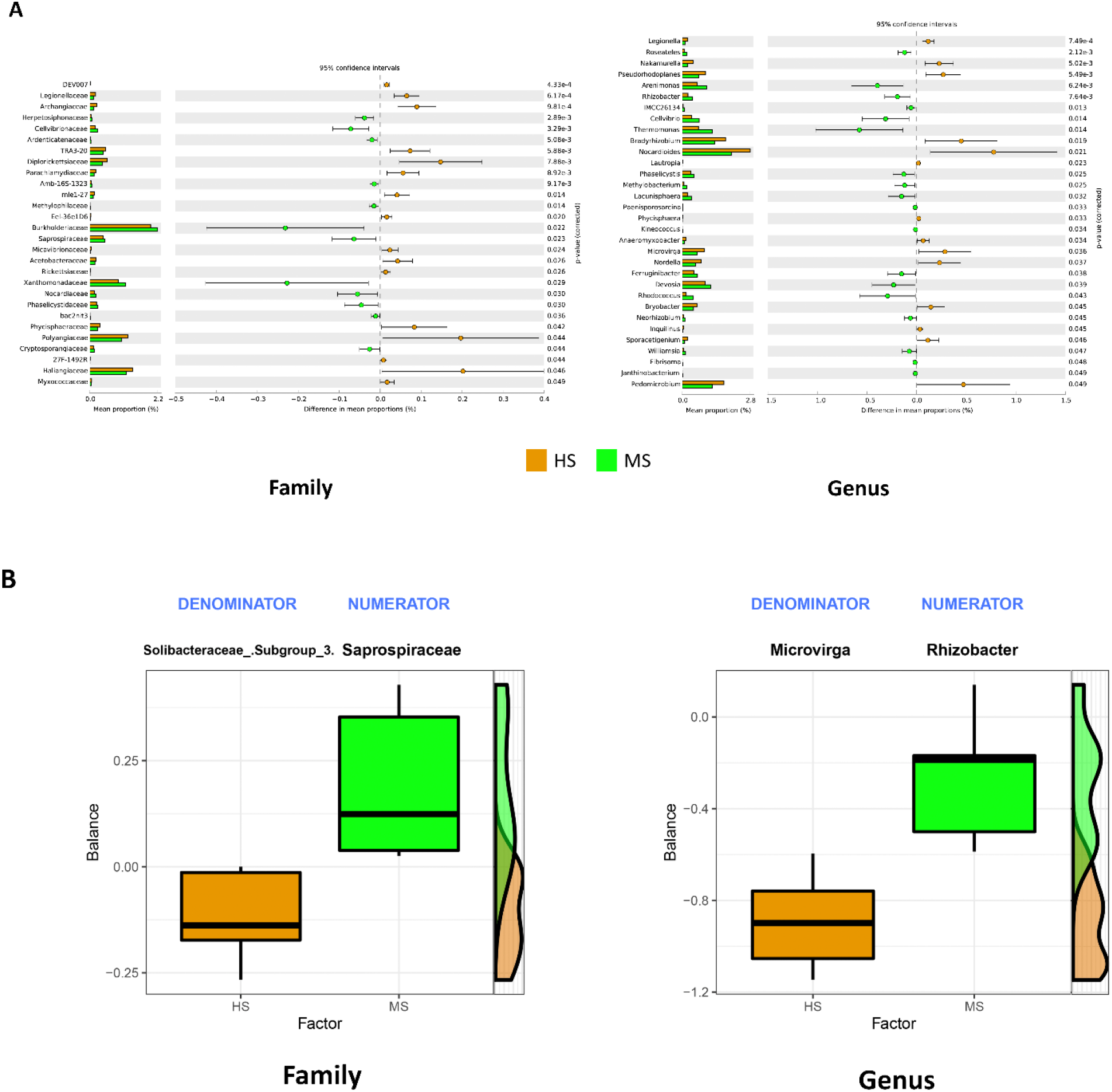
Rhizosphere microbial taxa relative abundance differences at family and genus level tested with Welch’s t-test (a). Compositional balance change analysis identifying the microbial signatures that discriminate the rhizosphere microbiome between HS and MS (b), the balance indicates the logarithm ratio of the relative abundance of identified denominator and numerator.

A co-occurrence network analysis was performed to visualize the microbial interaction of HS and MS turf rhizosphere soil bacteria and showed different network patterns (Fig. 7a). The co-occurrence networks were then further analyzed using “NetShift” to quantify the differences and identify the keystone microbial taxa that triggered the shift of the microbial networking between HS and MS rhizosphere bacterial communities when clustered at the Family and Genus level (Fig. 7b). There were 55 families and 28 genera identified as driver taxa when comparing HS and MS co-occurrence networks aggregated at each taxonomic level.

**Figure 7.**
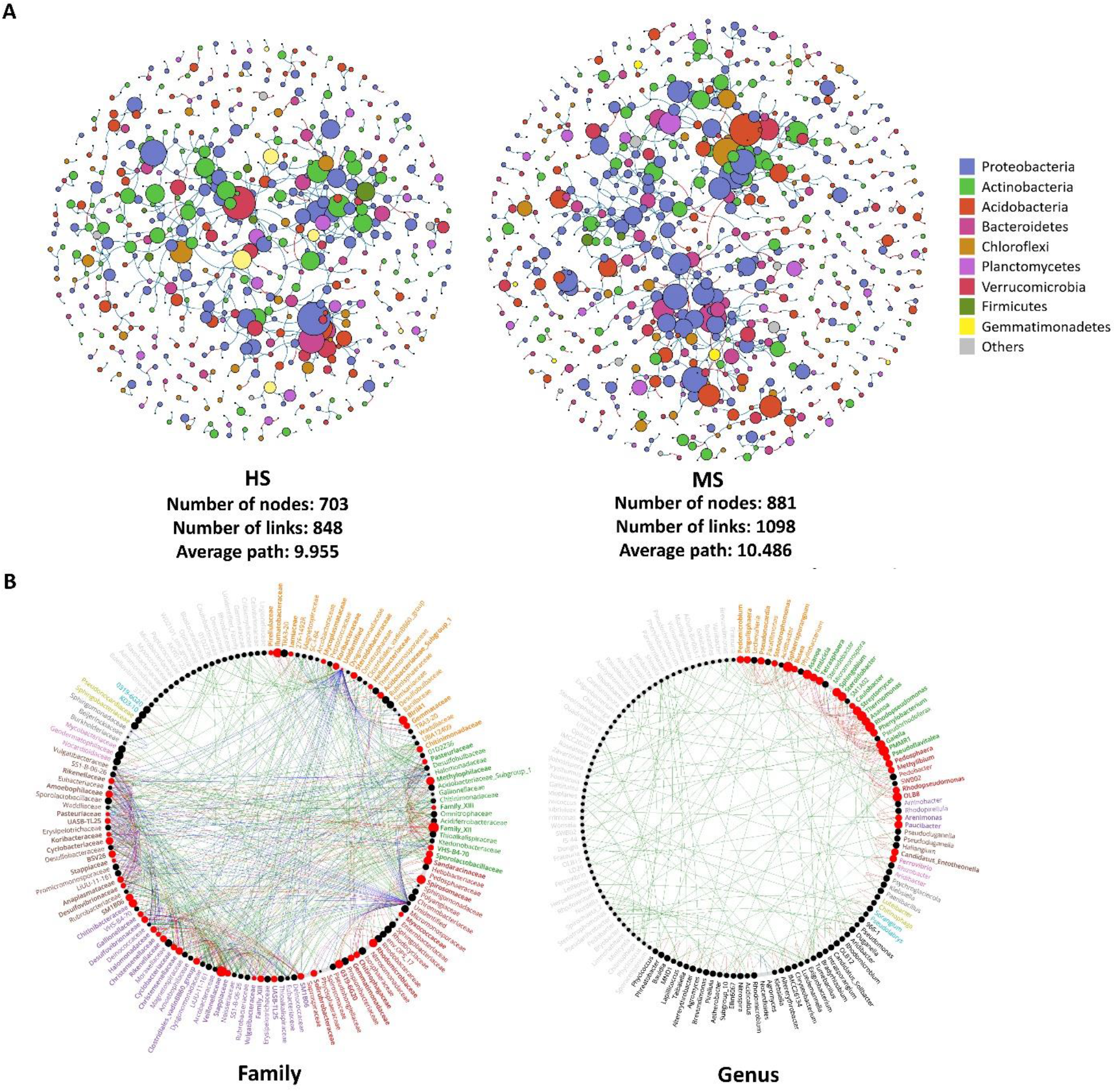
Rhizosphere soil bacterial microbiome co-occurrence networks at phylum level of dollar spot MS and HS turfgrass (a), in which the size of the nodes were scaled based on in-degree values, blue and pink paths represents positive and negative correlation, respectively. NetShift analysis by comparing the co-occurrence networks to identify the driver taxa at family and genus levels (b), where the nodes were scaled based on the degree in neighbor shift, the red nodes are the identified important drivers responsible for the network shift between the MS and HS turfgrass rhizosphere microbiome, and the green, red and blue paths represents the edges showed in MS, HS and both, respectively.

Rhizosphere soil bacterial function was predicted using Tax4Fun2 (21) to explore the potential microbial functional differences between HS and MS samples during the peak disease development period. Predicted functional pathways at level-two according to KEGG reference for molecular functions of genes (22) including nucleotide metabolism, folding, sorting and degradation, cell motility, translation, transcription, replication and repair, and metabolism of cofactors and vitamins associated genes were found to be more abundant in rhizosphere of MS samples (Fig 8). In the HS samples rhizosphere genes associated with xenobiotic biodegradation and metabolism pathways associated genes were more abundant (Fig. 8).

**Figure 8.**
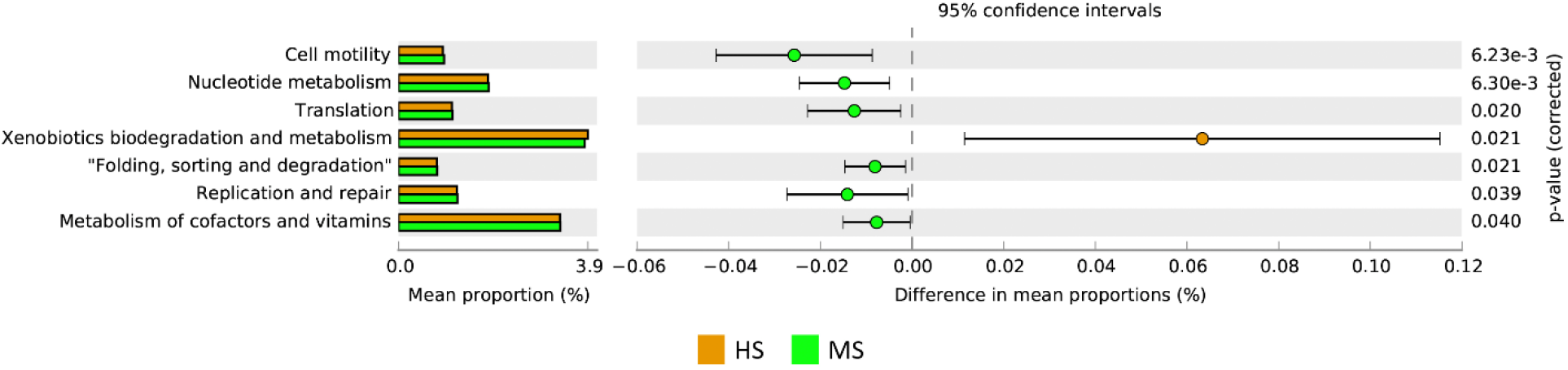
Significant differences in predicted rhizosphere microbiome functional pathways of MS and HS turfgrass using Tax4Fun2 and tested by Welch’s t-test.

### Bulk soil nutrient and chemical property analysis

Bulk soil chemical properties were compared among the three disease severity groups categorized according to turf dollar spot severity throughout the incubation period. The bulk soil was sampled prior to the inoculation of *C. jacksonii* to evaluate if bulk soil chemical property explained the turfgrass responses to the pathogen inoculation. The results showed that iron concentration was significantly lower in the high disease than the low disease group throughout the peak disease development stage from 4 to 10 DAI (Table 2), and iron was also lower in the HS samples relative to the MS samples (p=0.0021) following re-categorization of the samples (Table S1 in the supplemental material). A Mantel test was conducted to determine correlation between the overall soil chemical properties and the soil bacterial community. Bulk soil chemical properties did not correlate with the bulk soil bacterial community (r=−0.2297, p=0.966) but they did correlate with the rhizosphere bacterial community (r= 0.274, p=0.048). To further examine the relationship between bulk soil chemical properties and dollar spot severity during the peak disease development stage, a backward stepwise regression model was constructed after removing significant colinear variables. The stepwise model (adjusted r^2^=0.5041, p=0.002031) suggested that iron significantly (p=0.00062) and positively regressed with average turfgrass greenness during the peak development period (Table 3).

**Table 2.** Mean separation of turf-associated bulk soil chemical elements as categorized based on disease severity (high, medium, and low) after inoculation of *C. jacksonii* and throughout the incubation (a). Iron content (mg/kg of dry soil) of each severity group categorized based on the peak of disease development stage (b). Non-parametric Kruskal-Wallis test and Steel-Dwass paired-comparison were conducted to test the significance level. Asterix indicates the significance level: * p<0.05 and ** P<0.01. DAI stands for Days after inoculation of *C. jacksonii*. Numbers followed by ± indicates standard errors.

|  | DAI 0 |  | DAI 2 |  | DAI 4 |  | DAI 6 |  | DAI 8 |  | DAI 10 |  | DAI 12 |  | DAI 14 |  | DAI 16 |  |
| --- | --- | --- | --- | --- | --- | --- | --- | --- | --- | --- | --- | --- | --- | --- | --- | --- | --- | --- |
|  | ChiSq | P-value | ChiSq | P-value | ChiSq | P-value | ChiSq | P-value | ChiSq | P-value | ChiSq | P-value | ChiSq | P-value | ChiSq | P-value | ChiSq | P-value |
| pH | 0.626 | 0.060 | 4.526 | 0.104 | 3.170 | 0.205 | 1.509 | 0.470 | 1.064 | 0.587 | 2.561 | 0.278 | 2.012 | 0.366 | 2.667 | 0.264 | 2.667 | 0.264 |
| OM | 0.986 | 0.611 | 0.184 | 0.912 | 5.535 | 0.063 | 2.611 | 0.271 | 0.784 | 0.676 | 1.092 | 0.579 | 0.246 | 0.884 | 0.012 | 0.994 | 0.012 | 0.994 |
| Al | 3.942 | 0.139 | 6.398 | 0.041* | 8.082 | 0.018* | 5.661 | 0.059 | 4.012 | 0.135 | 9.310 | 0.01** | 3.310 | 0.191 | 3.193 | 0.203 | 0.319 | 0.203 |
| Ca | 1.977 | 0.372 | 2.854 | 0.240 | 5.719 | 0.057 | 9.310 | 0.01** | 2.561 | 0.278 | 5.485 | 0.064 | 1.450 | 0.484 | 0.889 | 0.641 | 0.889 | 0.641 |
| Cu | 2.117 | 0.347 | 10.714 | 0.005** | 4.667 | 0.097 | 2.328 | 0.312 | 0.363 | 0.834 | 2.538 | 0.281 | 3.170 | 0.205 | 0.924 | 0.630 | 0.924 | 0.630 |
| Fe | 5.099 | 0.078 | 3.509 | 0.173 | 9.275 | 0.01** | 7.906 | 0.019* | 8.924 | 0.012* | 11.614 | 0.003** | 4.714 | 0.095 | 4.994 | 0.082 | 4.994 | 0.082 |
| K | 0.363 | 0.834 | 0.152 | 0.927 | 3.521 | 0.172 | 7.029 | 0.030 | 3.193 | 0.203 | 2.047 | 0.359 | 0.328 | 0.849 | 1.263 | 0.532 | 1.263 | 0.532 |
| Mg | 1.063 | 0.587 | 2.538 | 0.281 | 3.661 | 0.160 | 4.678 | 0.096 | 0.854 | 0.653 | 3.170 | 0.205 | 1.275 | 0.529 | 0.667 | 0.717 | 0.667 | 0.717 |
| Mn | 3.029 | 0.220 | 0.877 | 0.645 | 1.205 | 0.548 | 1.509 | 0.470 | 2.632 | 0.268 | 3.895 | 0.143 | 0.503 | 0.778 | 0.246 | 0.884 | 0.246 | 0.884 |
| Mo | 0.456 | 0.796 | 0.222 | 0.895 | 0.714 | 0.700 | 5.556 | 0.062 | 1.556 | 0.459 | 0.714 | 0.700 | 0.105 | 0.949 | 0.737 | 0.692 | 0.737 | 0.692 |
| Na | 1.310 | 0.520 | 0.152 | 0.927 | 4.012 | 0.135 | 3.825 | 0.148 | 0.667 | 0.717 | 3.790 | 0.150 | 0.737 | 0.692 | 0.421 | 0.810 | 0.421 | 0.810 |
| P | 3.240 | 0.198 | 1.368 | 0.505 | 1.298 | 0.523 | 2.246 | 0.325 | 0.877 | 0.645 | 1.064 | 0.587 | 0.714 | 0.700 | 0.573 | 0.751 | 0.531 | 0.751 |
| S | 1.556 | 0.459 | 0.105 | 0.949 | 0.140 | 0.932 | 0.433 | 0.805 | 0.561 | 0.755 | 0.035 | 0.983 | 2.117 | 0.347 | 2.117 | 0.347 | 2.117 | 0.347 |
| Zn | 1.298 | 0.523 | 3.415 | 0.181 | 0.012 | 0.994 | 0.246 | 0.884 | 1.509 | 0.470 | 1.064 | 0.587 | 0.246 | 0.884 | 1.275 | 0.529 | 1.275 | 0.529 |
| C | 1.064 | 0.587 | 3.193 | 0.203 | 0.667 | 0.717 | 0.246 | 0.884 | 1.450 | 0.484 | 0.012 | 0.994 | 3.193 | 0.203 | 2.538 | 0.281 | 2.538 | 0.281 |
| N | 1.485 | 0.476 | 2.819 | 0.244 | 0.152 | 0.927 | 1.766 | 0.414 | 1.205 | 0.548 | 0.152 | 0.927 | 3.614 | 0.164 | 2.538 | 0.281 | 2.538 | 0.281 |

(B)
|  | Group | DAI 4 Mean |  |  |  | DAI 6 Mean |  |  |  | DAI 8 Mean |  |  |  | DAI 10 Mean |  |  |  |
| --- | --- | --- | --- | --- | --- | --- | --- | --- | --- | --- | --- | --- | --- | --- | --- | --- | --- |
| Fe | High | 0.825 | ± | 0.035 | b | 0.811 | ± | 0.037 | b | 0.818 | ± | 0.035 | b | 0.795 | ± | 0.031 | b |
| | Low | 0.989 | ± | 0.035 | a | 0.975 | ± | 0.037 | a | 0.987 | ± | 0.035 | a | 0.989 | ± | 0.031 | a |
| | Medium | 0.859 | ± | 0.035 | b | 0.887 | ± | 0.037 | ab | 0.868 | ± | 0.035 | ab | 0.889 | ± | 0.031 | ab |

**Table 3.**
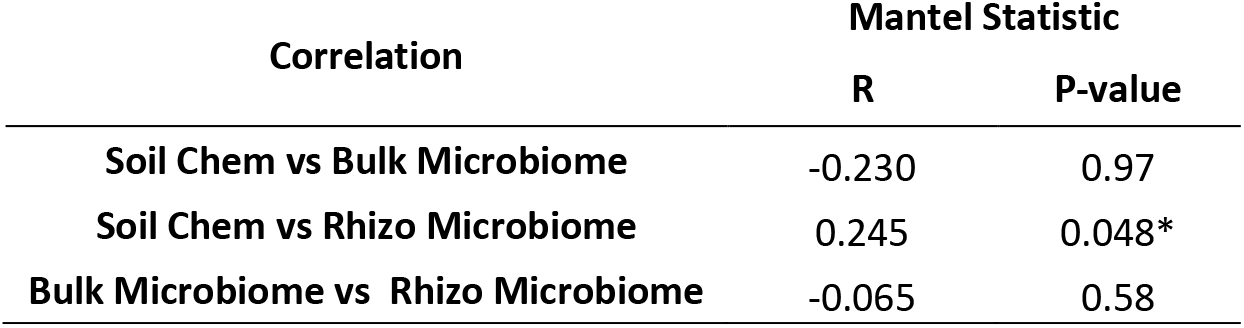
Correlations among bulk soil chemical property, bulk soil microbiome and rhizosphere microbiome using Mantel tests. Asterix indicates the significance level: * p<0.05.

**Table 4.**
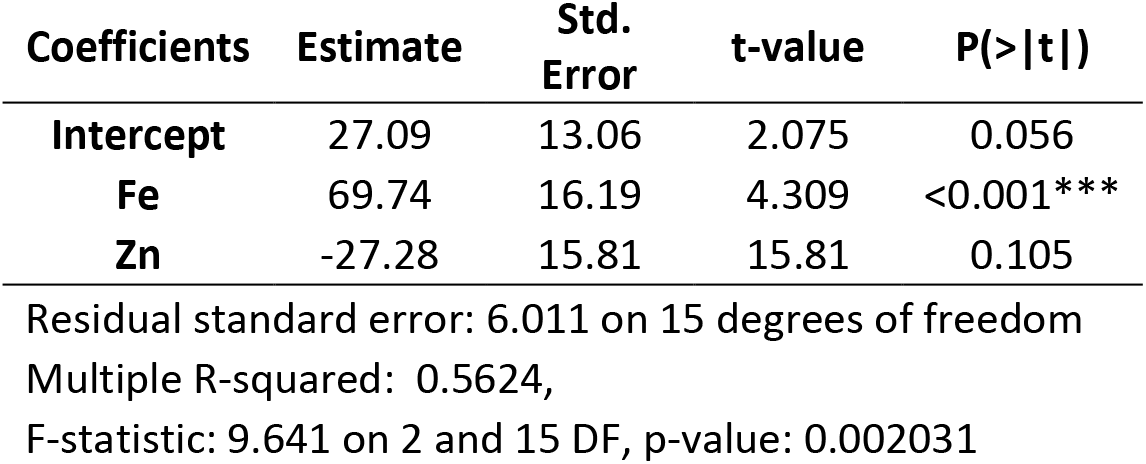
Stepwise selection of the optimal regression model for bulk soil chemical elements and average dollar spot disease severity (greenness) during the peak disease development stage (4-10 DAI). Asterix indicates the significance level: *** p<0.001.

## DISCUSSION

The results from this study indicated that initial differences in the soil rhizosphere bacterial community can predict the level of dollar spot susceptibility in turfgrass plants. These differences occurred over small areas despite uniform host plants and seemingly uniform environmental conditions. The mechanisms of disease suppression provided by the rhizosphere community were not directly studied, but differential analysis of microbial taxa relative abundances, and NetShift analysis of co-occurrence networks in this study provided supporting information for the hypothesis that disease suppression is related to the occurrence of antagonistic organisms in the rhizosphere. A similar hypothesis was also suggested in work done by Z. Wei et al. (19), which indicated that the rhizosphere bacterial community determined occurrence and severity of *Ralstonia solanacearum* in tomato plants and specifically linked disease suppression to the antagonistic activity of soil bacteria in the genera *Bacillus* and *Pseudomonas*. In our study, differential analysis revealed that certain families and genera were higher in relative abundance in the rhizosphere of MS samples compared to HS samples. These families, including *Nocardiaceae* and *Xanthomonadaceae*, and genera, including *Rhodococcus* and *Janthinobacterium*, are known to produce a range of antimicrobial compounds (23–26). Among the microbial co-occurrence network shift drivers identified through “NetShift”, node betweenness was significantly increased in MS samples for certain genera, including *Pseudonocardia*, *Streptomyces*, and “*Candidatus Entotheonella*,” which are all known for their ability to produce antifungal compounds (27–29). While more research is needed, these findings provide possible explanations for microbial suppression of dollar spot in MS turf samples.

In addition to known antibiotic producers, other bacterial taxa with environmental or plant functional importance in the rhizosphere differed between the HS and MS samples. The balance analysis revealed that the log ratios of *Saprospiraceae* and *Solibacteraceae* subgroup3 at the family level and *Rhizobacter* to *Microvirga* at the genus level can effectively differentiate between the rhizosphere microbiomes of the HS and MS groups. These results corresponded with differential relative abundance analysis as these taxa of microbial signatures were also captured by the differential relative abundance analyses. Microbial species under the genus *Microvirga* include many root symbionts (30), whereas members of the *Rhizobacter* genus are common rhizobacteria (31) and can also be plant pathogenic (32). Although the relative abundances were low, these identified taxa served as key signatures to differentiate the HS and MS rhizosphere bacterial community and may also have functional importance. For example, the identified family signature *Saprospiraceae* was present at a low level in our study (<1% in relative abundance), but members of the *Saprospiraceae* family are known to break down complex organic compounds in the environment (33) and are also suggested to have functional importance while underrepresented in soil abundance (34). The manner in which these microbial signatures interacted with the pathogen and host plant and whether they can be used for future evaluations of dollar spot suppression requires further research.

Functional prediction was performed to better understand implications of the differences identified in microbiome composition and interaction of HS and MS samples in the absence of a comprehensive metagenomic analysis. The MS rhizosphere microbiome was more enriched in genetic information processing and cellular processes metabolic pathways, whereas HS rhizosphere microbiome was more abundant in predicted xenobiotic biodegradation and metabolism. This result could help explain why the HS rhizosphere microbiome resulted in a more susceptible turfgrass sample. Many chemical compounds, such as salicylic acid (SA) analogs and β-Aminobutyric acid, can induce plant systemic acquired resistance that primes plants to defend against pathogens through activation of SA or abscisic acid (ABA) signaling pathways (35). Higher predicted abundance in gene associated with xenobiotic biodegradation and metabolism metabolic pathways in the HS rhizosphere microbiome suggested that the microbiome can more actively degrade xenobiotics such as agrochemicals, transformation products and secondary metabolites that either have direct antagonistic effects on pathogen growth, or compounds that have roles in priming plants against pathogens.

In the study by Z. Wei et al. (19), structural and functional differences in the rhizosphere microbiome were found to be the sole factors determining disease severity on tomato. In our study, bulk soil iron concentration predicted the disease susceptibility as well as that of the rhizosphere microbiome and seemed to contribute significantly to dollar spot suppression. S. Gu et al. (36) recently showed that siderophore production as a result of bacterial competition for iron resources in the soil environment strongly mediates *R. solanacearum* activity in the tomato rhizosphere. Specifically, iron-scavenging siderophores produced by nonpathogenic members of the bacterial consortia enhanced the fitness of these nonpathogenic bacteria in the soil environment and suppressed pathogen growth. Further large-scale screening of all major bacterial phylogenetic lineages established a strong positive linkage between inhibitory siderophore production by nonpathogenic bacteria and *R. solanacearum* suppression, indicating that the relative abundance of bacteria that produce pathogen-unusable siderophores in the tomato rhizosphere microbiome served as an effective predictor for disease outcome (37). These studies were done in a soil-borne pathosystem and it is unclear how pathogen-suppressing siderophore producers in the rhizosphere would compete with *C. jacksonii*, which is a foliar pathogen and poor soil saprophyte. Other mechanisms are likely involved, such as iron directly or indirectly neutralizing pathogen activity. For example, G. M. Gadd (38) observed that oxalic acid, a potential virulence factor of *C. jacksonii*, can react with the free iron in the plant-soil interface and precipitate as crystalline or amorphous solids. Also, in iron-deficient soils, induced bacterial production of the siderophore pyoverdine repressed the expression of plant defense-related genes such as the genes involved in SA and ABA pathways which can lead to a higher plant susceptibility to diseases (39).

Low soil iron can also lead to low iron in the plant tissue. Iron plays multifaceted roles in plant defense mechanisms and plant-pathogen interactions (40). For example, iron serves as a key factor in plant disease defense via numerous regulatory genes involved in microbe response and plant homeostasis, including upregulating the transcription of pathogenesis-related genes and catalyzing the reactive oxygen species when attacked by pathogens (41, 42). Unbalanced iron homeostasis in plants can have serious impacts on disease outcomes. Low iron in *Arabidopsis thaliana* led to more severe *Dickeya dadantii* infection due to less ferritin coding transcript AtFER1, callose deposition, and reactive oxygen species production (43). These collective studies on low soil and plant iron may help explain how lower soil iron in our study can lead to higher dollar spot susceptibility in turf and vice versa, but direct evidence on how soil iron interacts with the turfgrass plant to defend against dollar spot requires further analysis.

Numerous field and *in vitro* studies have shown the beneficial effect of iron in plant disease suppression (44–46), and the beneficial effects of iron are often found in conjunction with a pathogen-suppressive soil microbiome (14, 20). Healthy blueberry (*Vaccinium corymbosum*) plants were found to associate with more diverse rhizosphere bacterial communities and higher iron content in the roots compared with unhealthy plants (20). An *in vitro* study demonstrated that soil Fe-EDDHA amendment has an additive and complementary effect in suppressing Fusarium wilt (*Fusarium oxysporum* f. sp. *cubense*) disease severity in banana (*Musa spp*.) grown in a disease suppressive soil (14). The mechanisms of such a complementary effect of iron in our study remain unclear, but the Mantel test results suggest that the rhizosphere microbiome was likely mediated by interaction between soil iron levels and turfgrass plants, which in turn impacted disease development.

The rhizosphere microbiome is recruited or expelled from the bulk soil through the production of phytochemicals (47, 48) including many organic acids and secondary metabolites (49). More specifically, previous work by Y. Pii et al. (50) demonstrated that plant iron status had a significant impact on the formation of rhizosphere microbiome structures, possibly via the release of different qualitative and quantitative root exudates. In our study, higher Fe in the bulk soil of MS samples likely induced production of root exudates that then recruited a particular rhizosphere microbiome that was more suppressive to dollar spot development. However, this proposed mechanisms requires significant additional research before it can be used to develop innovative plant disease control strategies.

This study revealed several factors that led to variation in disease development over a small area in amenity turfgrass. Although further research is required before making firm conclusions, our findings suggest that antibiotic-producing members in the rhizosphere microbiome likely played a key role in the dollar spot suppression observed in MS samples. Further, soil iron-plant interactions were possibly a key regulatory factor in the assembly of a suppressive rhizosphere microbiome, and this soil-plant-microbe interaction ultimately resulted in the observed variation in disease development on monocultured turfgrass within a small scale. Future studies on whether the disease suppressive function can be transplanted into a conducive soil, and how turfgrass physiologically mediates root exudates to recruit a disease suppressive rhizosphere microbiome by responding to different levels of soil iron will be critical in further exploring the hypotheses raised by this research.

## MATERIALS AND METHODS

### Experimental design, sampling scheme and sample preparation

The experiment was conducted on a mature stand of creeping bentgrass (*Agrostis stolonifera* ‘Alpha’) at the O.J Noer Turfgrass Research Facility in Verona, WI, USA. The turf was grown on a native Troxel silt loam and mowed three times per week at the height of 1.25 cm. Eighteen turfgrass samples and the associated soil were taken using a soil sampler with a 13-cm diameter and a 15-cm depth in a 256 m^2^ square plot on Oct. 10^th^, 2019. The samples were divided into a top layer (the top 7.5 cm) and a bottom layer (7.5 to 15 cm depth) by carefully inserting the soil sampler to the specified depths. Due to the nature of the turfgrass and soil properties, there was hardly any soil without direct contact with roots in the top layer, and rarely root presence in the bottom layer soil. Therefore, we defined the bulk soil as the soil from the bottom layer without direct root contact. The soil samples of each layer were stored separately as turf and bulk soil samples. The turf samples were then used for inoculation experiments after they were sub-sampled for rhizosphere microbiome analysis. Bulk soil samples were sub-sampled from the homogenized bottom layer soil for both microbiome and chemical property analysis. Two, 1-cm diameter subsamples to 5-cm depth containing approximately 10 to 15 individual creeping bentgrass plants were taken from each turf sample for microbiome analysis using a custom-made soil probe. The subsamples from the same turf sample were immediately crushed with a sterile scapula and tweezer, and the soil loosely attached to the root system was separated from plant and rhizosphere soil by aggressively shaking in a sterile glass petri dish, rhizosphere soil remained closely attached to the root was then carefully collected using scapula avoiding the root tissues. The intact turf samples, which the subsamples were taken from, were then inoculated with one milliliter of dollar spot inoculum using a vaporizer within one hour of sampling. The dollar spot inoculum was created by growing *C. jacksonii on* potato dextrose broth for 72 hrs, rinsing three times in distilled water, and homogenizing in sterile 0.85% saline water in a blender for one minute. The final inoculum had an approximate *C. jacksonii* density of 4.1*10^4^ CFU/ml, as determined by testing with triplicated serial dilutions on potato dextrose agar.

After inoculation, the turf samples were incubated in a growth chamber at 25°C, 70% relative humidity, and 15 hr photoperiod. Each sample was placed on a sterile filter paper with an individual glass water pan. The turf samples were maintained at 0.5 cm height using sterile scissors, supplied with distilled water through wetting the filter paper, and measured for dollar spot severity every other day for 16 days (Fig. S1 in the supplemental material). Dollar spot severity was assessed by taking digital photos 30 cm directly above the turf surface and counting the percentage of green pixels using imageJ. Bulk soil samples were sent to the Cornell Nutrient Analysis Laboratory (Ithaca, NY) to analyze the chemical properties including pH, organic matter content, and Al, Ca, Cu, Fe, K, Mg, Mn, Mo, Na, P, S, Zn, C and N content according to procedures outline in B. K. Gugino et al. (51). Briefly, soil were dried in open containers overnight, sieved to remove pebbles and plant tissues, soil organic matter content was measured by dry combustion at 550°C for two hours, and pH was measured as 1:1 soil to water solution by volume using an automatic pH probe (Lignin, Albuquerque, NM). Soil nutrients were extracted using Morgan’s solution and quantified with an Inductively Coupled Argon Plasma Spectrophotometry (Thermo Fisher Scientific, Cambridge, UK).

### Library Preparation and short-amplicon sequencing

For each of the bulk soil and rhizosphere soil samples, 0.25 g soil was used for DNA extraction using a DNeasy PowerLyzer PowerSoil kit (Qiagen Inc., Germantown, MD) following the manufacturer’s protocol. All extractions were quantified for nucleic acid concentration using a NanoDrop1000 (Thermo Fischer Scientific, Waltham, MA). The PCR was performed according to K. A. Dill-McFarland et al. (52) with minor modifications. Briefly, each reaction contained 5 μL of the DNA template at 10 ng/μL, 12.5 μL Kapa HiFi HotStart ReadyMix, 6.5 μL PCR-grade water, and 0.5 μL of each barcoded forward and reverse primer (53), which targeted the v4 region of the 16S rRNA gene. The thermocycling conditions were 3 min at 95 °C prior to 25 cycles of 30 s at 95 °C, 30 s at 55 °C, and 30 s at 72 °C, with a final step of 5 min at 72 °C. The amplicons were purified using a ZR-96 Zymoclean™ Gel DNA Recovery kit (Zymo Research, Irvine, CA) and normalized with a Mag-Bind^®^ EquiPure Library Normalization Kit (Omega Bio-Tek Inc, Norcross, GA). The amplicons were then pooled and quantified to 4 nM with a Qubit™ dsDNA HS Assay kit (Thermo Fischer Scientific, Waltham, MA). The final pool was sequenced on Illumina MiSeq with a 2×250bp PE Illumina Reagent Kit v2 (Illumina, Inc., San Diego, CA) in the Biotechnology Center at the University of Wisconsin-Madison

### Data analysis

The raw sequences were processed using package “DADA2” in R 3.6.0. Forward and reverse reads were quality filtered according to average quality score and merged. The taxonomy levels associated with each amplicon sequence variant (ASV) was assigned according to SILVA database (v.132) after removing the chimeras. The ASV and taxonomic tables were then exported as .txt files and analyzed using R packages “phyloseq” and “vegan.” The reads for each sample were normalized using variance stabilizing transformation with the “DeSeq2” package due to a relatively even reads variation among the samples in the library (54). Microbial compositional differences and correlations were analyzed using Bray-Curtis dissimilarity. Shannon diversity of HS and MS were compared using nonparametric Wilcoxon test in JMP Pro 14 (SAS Institute, Cary, NC).

Microbial co-occurrence network of HS and MS samples were constructed using Molecular Ecological Network Analysis (MENA) (55), which uses a Random Matrix Theory (RMT)-based method to predict the microbial interactions and capture the magnitude of the interactions. The nodes and the edge lists were then imported into Gephi 0.9.2 (56) for network visualization. Since the overall ASVs were comprised of approximatly 90% of the ASVs having less than 0.02% of overall reads, ASVs that represent less than 0.02% of the total reads after normalization for each sample were filtered out to make the result more readable. The core community of the HS and MS microbial networks were compared to quantify the rewiring of the taxa in the networks by calculating the of neighborhood shift and change of betweenness for the nodes using NetShift (57). Nodes with the highest degree change among these parameters are considered the driver taxa. When analyzed at family and genus level, the ASVs were aggregated at each taxonomy level to create the edge list. Microbial balance analysis was performed using “selbal” package in R at family and genus level using unnormalized ASV counts, as the compositional nature of the short-amplicon sequencing result and the uneven sequencing depths were both accounted in the analysis (58). Differential relative abundances were analyzed using Welch’s t-test at a significance level of α=0.05 in using Statistical Analysis of Taxonomic and Functional Profiles (STAMP) (59).

Rhizosphere microbiome functional prediction was performed using an R-based tool Tax4Fun2 (21), which used the sequences of the ASV to blast against the SILVA (v.132) reference genome database to create a metagenome profile. The genetic functions were then assigned by BLASTp against the KEGG KO (22) as a reference database. Differences in functional pathways at level-two were statistically analyzed using Welch’s t-test in STAMP. The associations of Bray-Curtis dissimilarity among bulk soil chemical properties, bulk soil microbiome, and rhizosphere microbiome samples were examined using Mantel test in R.

Soil chemical properties among the disease groups were statistically analyzed with nonparametric Wilcoxon test in JMP Pro 14 (SAS Institute, Cary, NC) and regression with average disease severity of peak disease development stage (4-10 DAI) was performed using a stepwise selection for the optimal predictive model in R. Collinearity variable selection and removal was performed using a customized function vif_func (60) to calculate the variance inflation factor. The best model was constructed with backward selection using a function stepAIC under package “MASS”.

### Data availability

All the raw sequences generated from this study were deposited at the NCBI Sequence Read Archive and are publicly accessible under the project number of PRJNA642971.

## ACKNOWLEDGEMENT

The authors thank Kurt Hockmeyer and Qiwei Lei for assistance in sampling, Joseph Skrlupka and Dr. Garret Suen for assistance in preparation for sequencing, and the UW-Madison CALS Statistical Consulting Lab for providing guidance in statistical analysis.

---

**Supplementary Table S1.** Wilcoxon non-parametric comparison of soil chemical properties between HS and MS associated bulk soil.

| Property | Level | Mean | Std. Dev. | Std. Error<br>Mean | Lower<br>95% | Upper<br>95% | Wilcoxon p-value |
| --- | --- | --- | --- | --- | --- | --- | --- |
| pH | HS | 7.240778 | 0.233394 | 0.0777981 | 7.061375 | 7.420181 | 0.0543 |
|  | MS | 7.398875 | 0.07535 | 0.0266401 | 7.335881 | 7.461869 |  |
| OM | HS | 2.985949 | 0.275381 | 0.0917936 | 2.774273 | 3.197625 | 0.7361 |
|  | MS | 3.076462 | 0.103921 | 0.0367415 | 2.989582 | 3.163342 |  |
| Al | HS | 2.774205 | 0.252402 | 0.0841339 | 2.580192 | 2.968218 | 0.0161* |
|  | MS | 3.162284 | 0.29524 | 0.104383 | 2.915458 | 3.409111 |  |
| Ca | HS | 1639.915 | 80.10273 | 26.700911 | 1578.342 | 1701.487 | 0.0433* |
|  | MS | 1714.069 | 110.1071 | 38.928753 | 1622.017 | 1806.121 |  |
| Cu | HS | 0.080371 | 0.032808 | 0.0109359 | 0.055153 | 0.105589 | 0.2482 |
|  | MS | 0.092079 | 0.020072 | 0.0070966 | 0.075298 | 0.10886 |  |
| Fe | HS | 0.818683 | 0.097546 | 0.0325154 | 0.743702 | 0.893664 | 0.0021** |
|  | MS | 0.975093 | 0.055358 | 0.0195718 | 0.928813 | 1.021373 |  |
| K | HS | 155.3454 | 11.78698 | 3.9289926 | 146.2852 | 164.4057 | 0.1489 |
|  | MS | 161.5223 | 8.360104 | 2.955743 | 154.5331 | 168.5115 |  |
| Mg | HS | 493.3613 | 26.29071 | 8.7635684 | 473.1525 | 513.5702 | 0.2898 |
|  | MS | 507.6362 | 38.32882 | 13.551285 | 475.5925 | 539.6799 |  |
| Mn | HS | 2.186369 | 1.367373 | 0.4557909 | 1.135313 | 3.237424 | 0.0833 |
|  | MS | 3.398221 | 1.248337 | 0.4413539 | 2.354585 | 4.441857 |  |
| Mo | HS | 0.005837 | 0.001988 | 0.0006626 | 0.004309 | 0.007364 | 0.9233 |
|  | MS | 0.005627 | 0.001894 | 0.0006696 | 0.004044 | 0.00721 |  |
| Na | HS | 27.50916 | 2.103361 | 0.7011202 | 25.89237 | 29.12594 | 0.5006 |
|  | MS | 28.36482 | 1.818714 | 0.6430124 | 26.84434 | 29.8853 |  |
| P | HS | 16.69933 | 1.557548 | 0.5191827 | 15.50209 | 17.89657 | 0.4414 |
|  | MS | 15.66392 | 2.09468 | 0.7405812 | 13.91272 | 17.41511 |  |
| S | HS | 4.721017 | 0.446798 | 0.1489327 | 4.377577 | 5.064456 | 0.5006 |
|  | MS | 4.889829 | 0.443375 | 0.1567568 | 4.519157 | 5.2605 |  |
| Zn | HS | 0.825432 | 0.158275 | 0.0527584 | 0.703771 | 0.947093 | 0.8474 |
|  | MS | 0.849113 | 0.034658 | 0.0122533 | 0.820139 | 0.878088 |  |
| C | HS | 1.798 | 0.085772 | 0.0285905 | 1.73207 | 1.86393 | 0.9233 |
|  | MS | 1.843875 | 0.214384 | 0.0757961 | 1.664646 | 2.023104 |  |
| N | HS | 0.170222 | 0.011998 | 0.0039992 | 0.161 | 0.179445 | 0.8474 |
|  | MS | 0.176125 | 0.021649 | 0.0076542 | 0.158026 | 0.194224 |  |

**Supplementary Figure S1.**
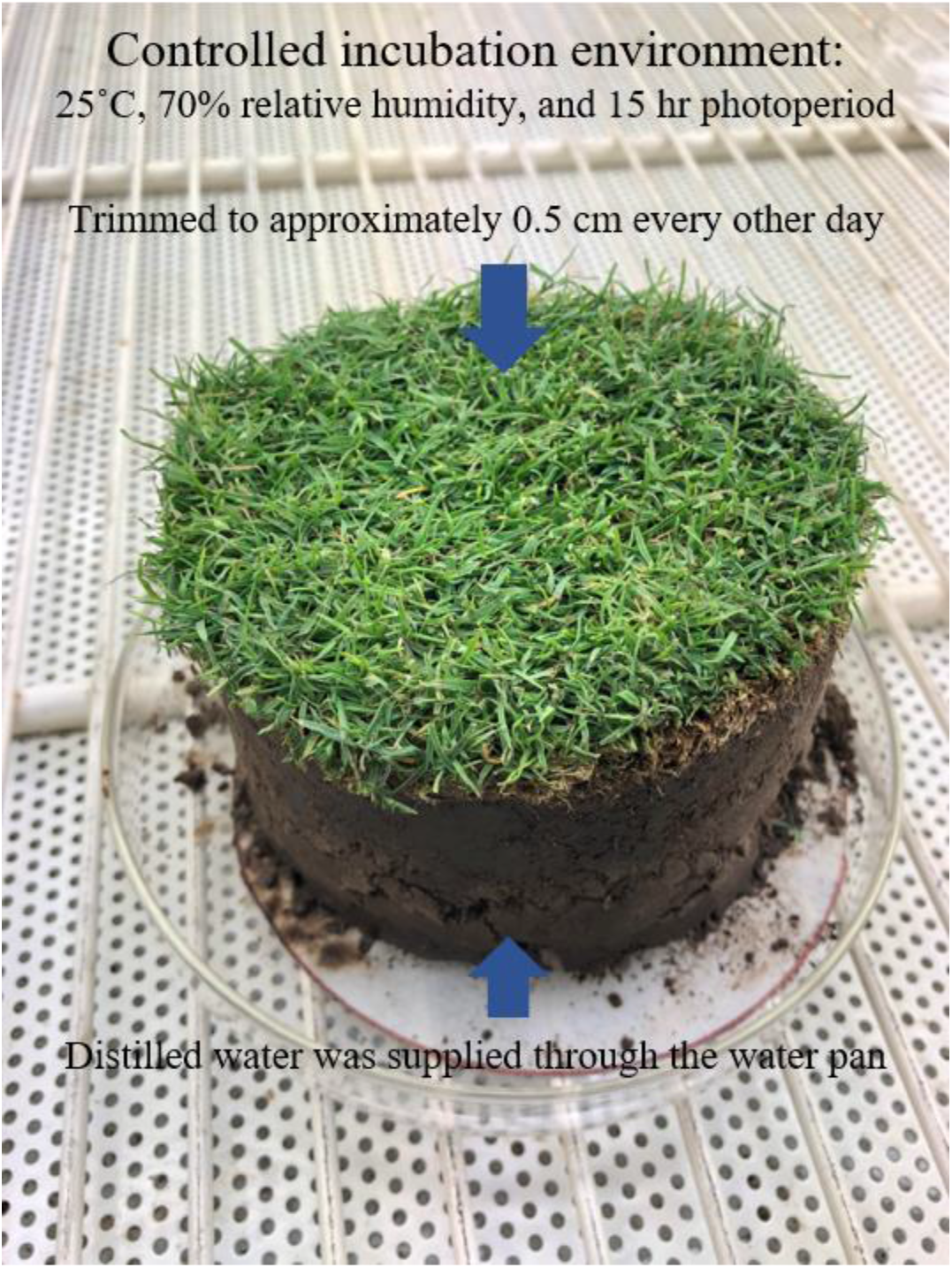
Set-up of each turf sample in the controlled environment growth chamber for the incubation after inoculation with *C. jacksonii*.

